# Unintended Consequences and the Paradox of Control: Management of Emerging Pathogens with Age-Specific Virulence

**DOI:** 10.1101/149773

**Authors:** Spencer Carran, Matthew Ferrari, Timothy Reluga

## Abstract

We project the long term incidence of Zika virus disease (ZVD) under varying hazards of infection and consider how the age-distribution of disease burden varies between these scenarios. Pathogens with age-structured disease outcomes, such as rubella and Zika virus, require that management decisions consider their impact not only on total disease incidence but also on distribution of disease burden within a population. In some cases, reductions of overall transmission can have the paradoxical effect of increasing the incidence of severe disease despite decreasing the total incidence. This happens because of corresponding increases in the average age of infection. Beginning with the current population structure and demographic rates of Brazil, we project forward total ZVD burden as measured by cases occurring in pregnant women and document the scenarios under which a paradox of control for Zika management emerges. We conclude that while a paradox of control can occur for ZVD, the higher total costs from increasing the average age of infection will only be realized after several decades and vanish under conservative discounting of future costs. This indicates that managers faced with an emerging pathogen should prioritize current disease incidence over potential increases in severe disease outcomes in the endemic state.

**Author Summary:** The intuitive response to an emerging outbreak is to halt, or at least reduce, transmission. However, in some circumstances, reducing overall transmission and incidence may be counterproductive from a public health perspective as public health interventions affect both the total level and the distribution of disease burden. We consider the scenarios under which reducing transmission of an emerging pathogen such as Zika virus may increase the costs associated with disease in the most vulnerable segments of the population - in this case, reproductive-age women. We conclude that after applying standard discounting rates to future cases, the “paradox of control” vanishes and reducing hazard of infection uniformly reduces the total costs associated with severe disease.

## Introduction

Zika virus (ZIKV), a mosquito-borne pathogen first identified in Uganda in 1947 [1] and subsequently responsible for sporadic outbreaks [2], has attracted major attention from health officials and the public at large as a result of an ongoing large outbreak in the Americas. The American Zika virus disease (ZVD) outbreak began in Brazil and rapidly spread through South and Central America, with an estimated 500,000-1,500,00 cases in Brazil alone [3]. While Zika virus disease (ZVD) is usually asymptomatic or mild [2], it has been linked to more severe complications in pregnant women [4]. The complication of greatest concern is microcephaly, where ZVD infection during fetal development impedes brain development.

Concerns over microcephaly have led to calls for women to delay or strategically time pregnancy [5, 6]. However, given the limited access to contraception and family planning services in much of Latin America [7], it may be more practical to focus on population-level control efforts that do not rely on individual behavioral modification. In particular, there has been renewed attention on potential vector control strategies [8] to reduce the attack rate (fraction of susceptible individuals experiencing infection) across the entire population.

For many diseases, minimizing attack rate is a straightforward way to reduce disease-associated mortality and morbidity. Attack rates determine not only the overall level of incidence, but also the average age of infection, with higher attack rates resulting in lower average age of infection [9]. For a disease which causes the most severe outcomes in younger individuals, such as measles, this suggests that reducing incidence also shifts the burden of disease away from the most vulnerable individuals. For a disease in which outcome severity can increase with age, such as rubella [10–12] or ZVD, decreases in the attack rate can shift cases into more vulnerable age classes. This may result in a”paradox of control” in which a reduction in incidence increases mortality and morbidity [13, 14]. The paradox of control can lead to situations with multiple locally optimal management equilibria [15, 16] and is the reason current WHO policy for rubella vaccination does not recommend implementing routine coverage below a threshold that is expected to reduce both total incidence and incidence in most-affected classes. The tradeoff in (usually mild) cases averted to a potential increase in incidence of congenital rubella syndrome (CRS) as seen in the Greek experience [17] is deemed unacceptable.

Responses to ZIKV spread should consider the trade-off rubella responses face with CRS. Otherwise, efforts to minimize near-term ZDV incidence may inadvertently shift the burden of disease to more vulnerable subpopulations. However, analyses of the tradeoff between rubella incidence and CRS burden [13, 18] have been based on an equilibrium incidence assumption. As ZVD is a newly emerging pathogen in a previously naïve population, it is not at equilibrium yet and we do not know what the incidence and age distribution will be at equilibrium. Additionally, our understanding of ZVD is rapidly expanding as new control methods such as genetically modified mosquitoes [19] and ZIKV vaccines [20–22] may change the eventual equilibrium level and distribution of ZIKV incidence.

We believe that cost-benefit evaluations of ZIKV policy interventions should focus primarily on the transient dynamics with discounting of future cases as is common in the economic literature [23]. In this paper, we use an age-structured model to study the potential short-term and long-term consequences of changes to a constant background ZIKV attack rate on incidence of ZVD and of high-risk cases in reproductive-age women during the transient dynamics following introduction. Our results show that the paradox of control is much weaker under transient dynamics, and almost always vanishes under even conservative discounting rates. We conclude that early interventions that reduce attack rates will always improve public health, and the paradox of control need only be considered when interventions have been delayed to a time when incidence has approached equilibrium levels.

## Materials and Methods

To evaluate the impact of an emerging pathogen with age-structured virulence, we construct a two-part model characterizing the underlying demographic structure, which can be well described with available census data, and overlay a disease incidence model, which describes a process with greater uncertainty. For our case we consider the initial conditions as the population structure of Brazil in 2015 (Fig 1) since Brazil was the most heavily impacted country in the recent outbreak and presents an interesting case study given a current age distribution that is disproportionately skewed towards the most vulnerable age classes.

**Fig 1.**
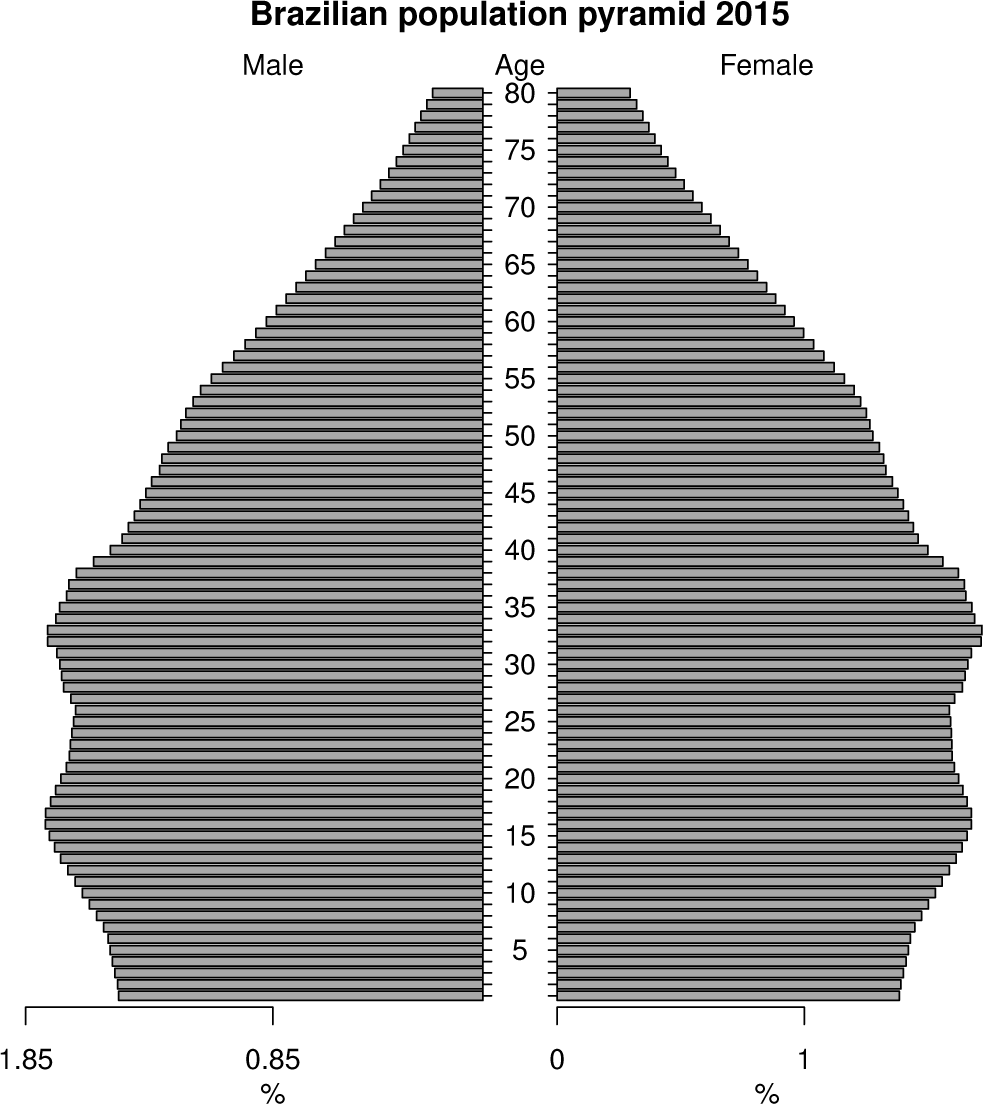
Brazilian Population Pyramid, 2015. Brazil’s population currently has the bulk of its mass in reproductive-age individuals, complicating any recommendation to delay childbearing.

The population is projected forward using recently estimated age specific fertility rates [24] (Fig 2). We assume individuals are born susceptible and removed from the susceptible population at an annual rate corresponding to the hazard of ZVD, and that individuals who are infected once retain lifetime immunity to future infection. We then consider the burden of ZVD in terms of the risk of ZVD-related birth defects. Actual rates of ZVD-related birth defects in different settings have been estimated as ranging from 11% [25] to 42% [26] and may vary by stage of pregnancy [27]. For purposes of our model, the total cost of ZIKV is defined as the number of births that occur in women who experience ZVD in the same year as their pregnancy.

**Fig 2.**
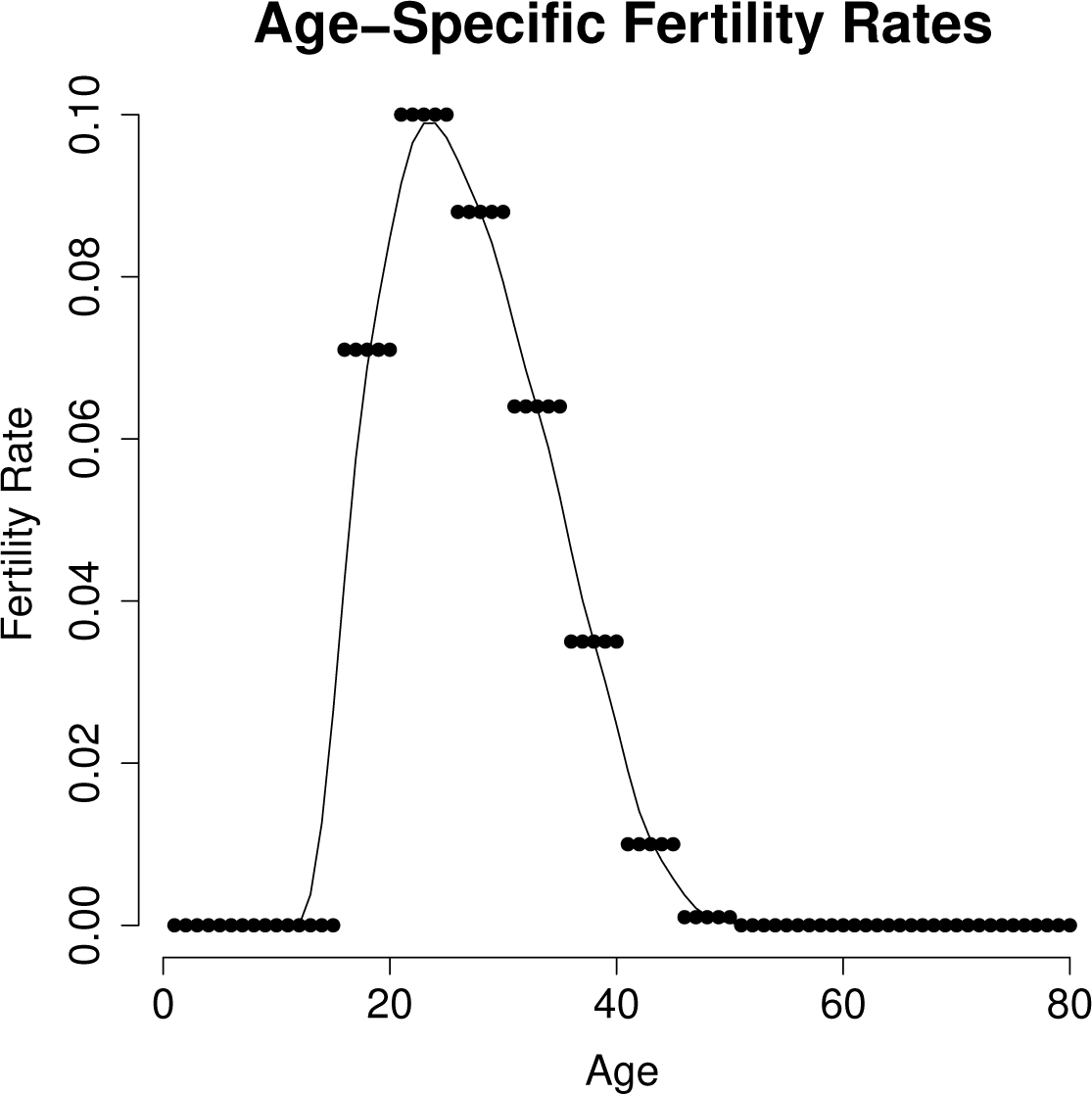
Age-Specific Fertility Rates. Age-specific Brazilian fertility rates as of 2012 in five-year intervals, with a smoothing spline fit to obtain annual resolution.

We project forward population dynamics and ZVD incidence over fifty years to generate a cumulative cost of ZIKV. In light of potential improvements in prenatal care for pregnancies coinciding with ZVD or control methods such as the release of genetically modified mosquitoes (citation) or a vaccine (citation) we weight present cases more heavily than future cases using the geometric rate 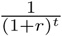 for cases *t* years in the future and an annual discounting rate *r*. Such discounting is standard practice in many areas of social policy [23].

Given that the hazard of encountering ZIKV infection is uncertain and likely to vary both in time and across spatial scales, we generate projections across a wide range of potential hazard rates. We consider the cumulative costs of ZIKV as a function of hazard rates in order to identify the possibility of a paradox of control under the assumption that ZVD hazard, while unknown, may be increased or decreased as a function of the intensity of control efforts.

Our model is a numerical approximation of an age-structured epidemic model with time-dependent infection risks, combined with a Lotka’s boundary condition for birth. Models with similar forms have been studied since the 1920’s [28, 29], based on McKendrick’s partial differential equation [30]. Let *S*(*t*, *a*) be the density of susceptible individuals of age *a* at time *t*. We assume a perfect sex ratio of 50/50. Individuals die at rate *μ*(*a*), depending on their age, and become infected at rate *λ*, independent of age and time. Infected individuals are assumed to become permanently immune against infection as soon as they are infected. New susceptible individuals are born at rate l(a) per susceptible person of age *a.* We assume ZVD infection has no measurable impact on the population’s large-scale demographic structure, so *l*(*a*) can be picked to reflect the collective birth rate of susceptible and resistant individuals. Thus
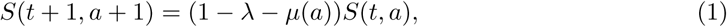

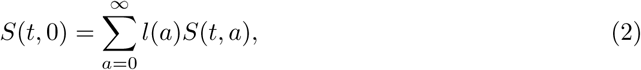

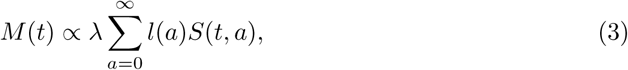

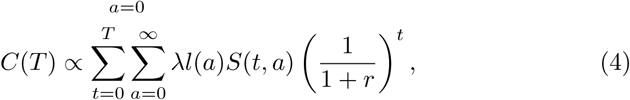

with the initial age-distribution of susceptibles *S*(0, *a*) and the maternity function *l*(*a*) estimated from census data [24].

The annual number of at-risk births *M*(*t*) in year *t* is proportional to the infection hazard *λ* and the total number of susceptible births. The cumulative discounted future cost of the Zika epidemic *C*(*T*) is proportional to the total number of at-risk births from the start of the epidemic up until year *T*, discounted at annual rate *r*.

Since the actual hazard of ZVD is uncertain, we consider the incidence of ZVD in high-risk age classes under varying annual hazard rates of ZIKV infection. We compare both the year-over-year and cumulative incidence of ZVD in at-risk age classes over a fifty year time window, and consider the impact of discounting future costs at the geometric rate 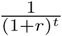 for cases *t* years in the future and a discounting rate *r*. Such discounting is standard practice in many areas of social policy [23]. Changes in the annual hazard rate (potentially modulated by control intensity) result in changes to both the equilibrium incidence and average age of infection. We have also explored some cases of age and time-dependent infections hazards (λ(*a*, *t*)), notably oscillating hazard rates across different years (supplement) and find no impact on our qualitative conclusions.

## Results

Considering the potential total number of at-risk births over the duration of 50 years with varying levels of ZVD incidence yields projections where intermediate levels of ZVD incidence lead to the highest total number of at-risk births while extremely high or extremely low ZVD incidence both result in a lower total burden. Since ZVD is an emerging infection, the age distribution of cases following introduction will simply match the population’s age distribution. As ZVD becomes established in a population, the age distribution will begin to shift towards younger individuals [9, 31]. The effect of the shifting age distribution is seen as the cost of ZVD as measured in cases in pregnant women tends to decline for any given hazard rate until an equilibrium is reached (Fig 3).

**Fig 3.**
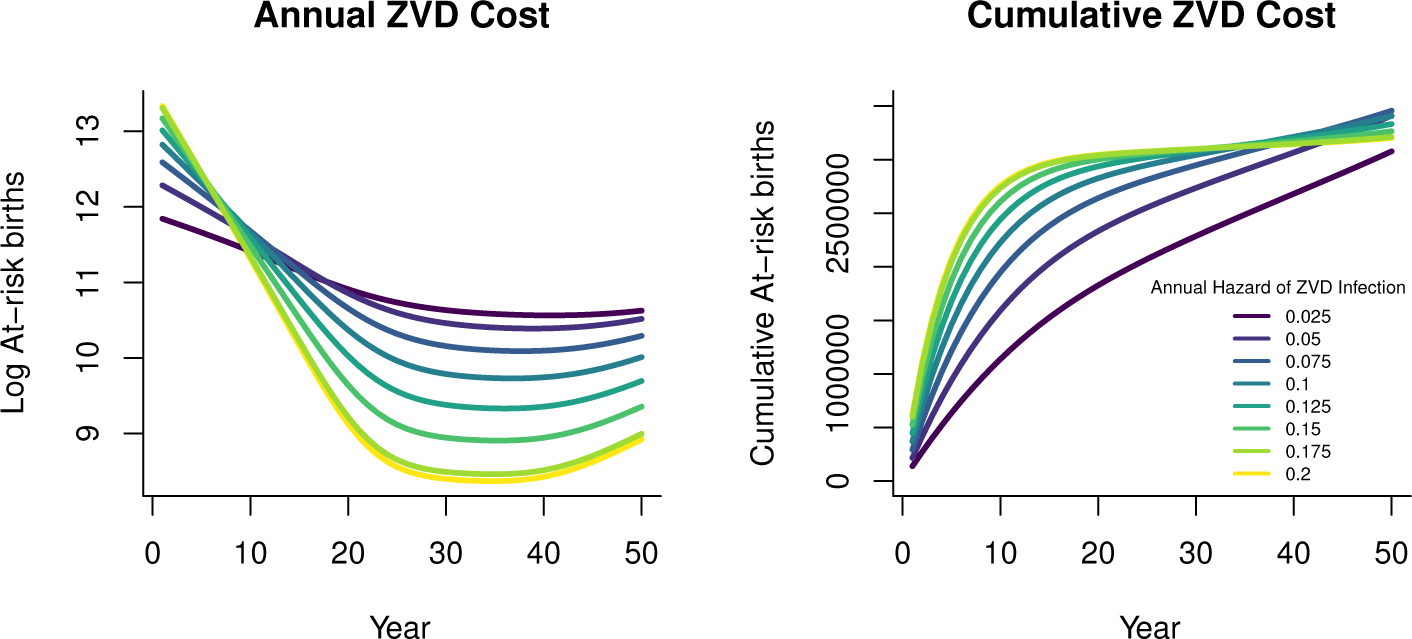
Annual (left) and cumulative (right) projected number of at-risk births over time in a population matching Brazilian age structure and demographic rates. A larger ZVD hazard results in more at-risk births at the start of the outbreak, while fewer people would remain susceptible in the long term. Scenarios involving lower ZVD hazard lead to fewer cumulative at-risk births over short planning horizons but eventually exceed high-hazard scenarios.

The cumulative burden of ZVD and microcephaly is determined both by the transient spread of ZVD through the initially naive population and the long-term endemic level of incidence once ZVD is established within the population. While high hazard rates lead to a larger initial outbreak, they also result in most of the population acquiring immunity before reaching reproductive age, limiting the potential for microcephaly in the future.

Placing possible hazard rates on the x axis, we consider the total cost over a fifty year window (Fig 4). Under the parameters we used, the greatest total burden of ZVD occurs when the annual hazard of contracting ZVD is 0.09, implying that efforts to reduce transmission in regions where hazard is higher than that could be counterproductive unless they succeed in reducing hazard below that threshold. For example, reducing the annual hazard from 0.15 to 0.12 would result in an estimated 36,500 additional cases of ZVD among pregnant women during the fifty year window of our projection.

**Fig 4.**
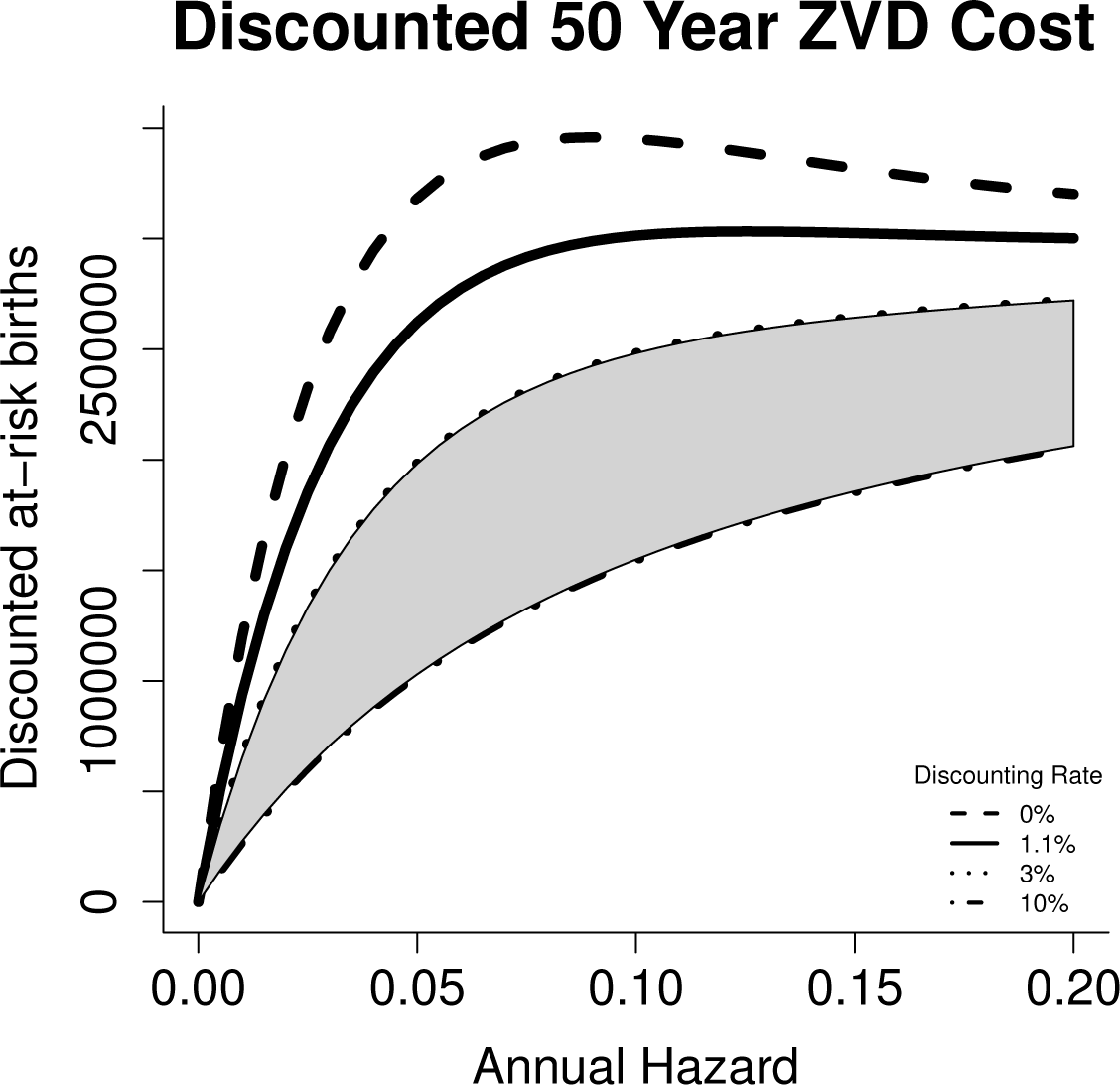
Intermediate levels of ZVD hazard lead to the largest number of at-risk births. When annual discounting of 1.1% or greater is applied to future cases, the total weighted cost of ZVD increases monotonically with annual hazard. The gray region indicates the range of discounting rates commonly used in social policy [23].

**Fig 5.**
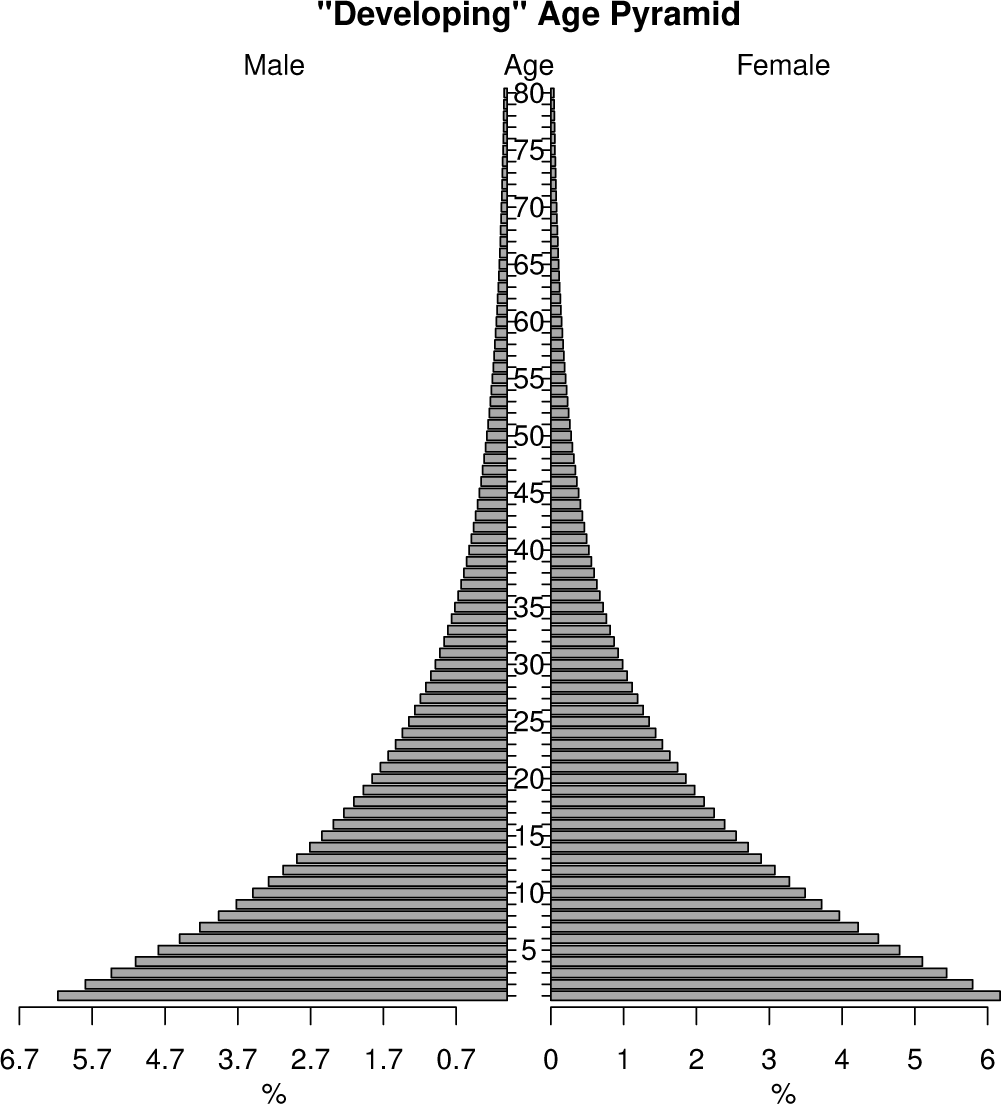
“Developing” Age Pyramid. In a setting with far more children (proportionally) there is less cost, and more potential benefit, to higher ZVD hazards providing a “natural vaccine” to ZVD infection during pregnancy

**Fig 6.**
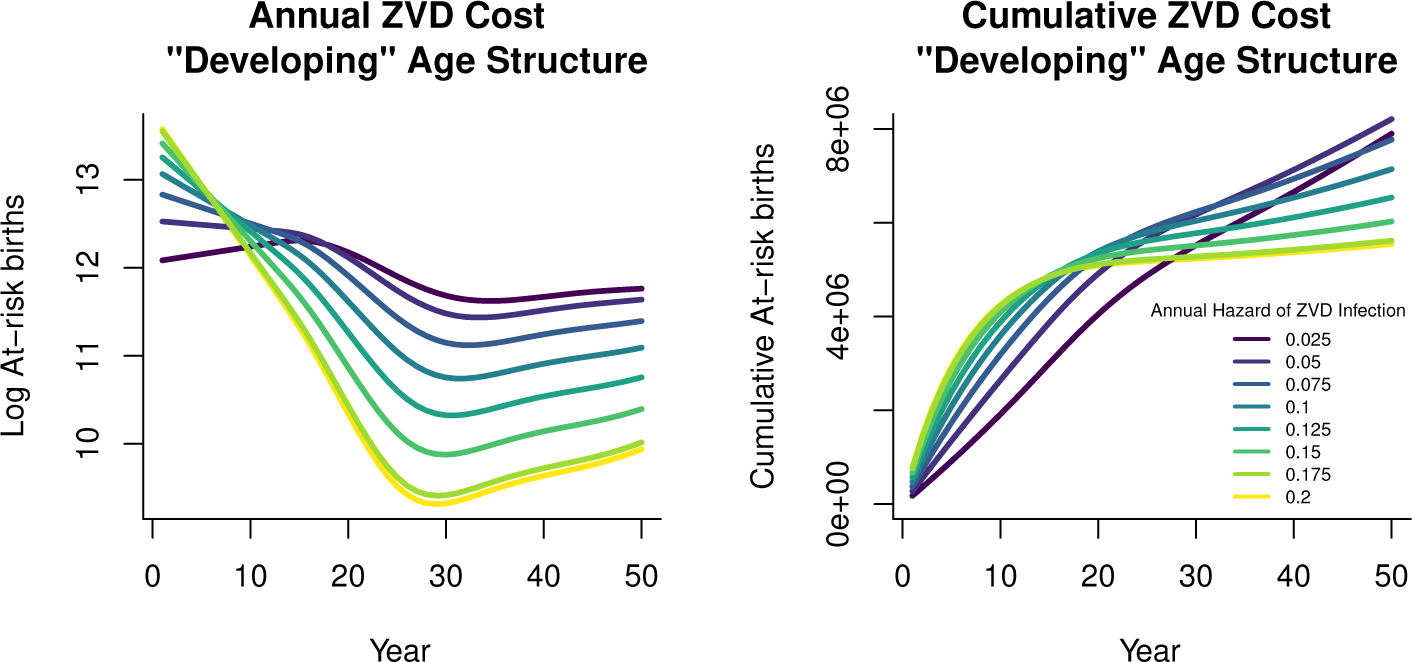
Since most individuals in this hypothetical developing population (see Figure 5) are below reproductive age, costs of ZVD tend to be higher in the future than in the case of Brazilian demography. The left panel shows absolute number of annual ZVD cases in pregnant women under varying hazard rates, while the right panel shows cumulative number of cases. This assumes the starting population age distribution is exponential and begins with all individuals susceptible.

**Fig 7.**
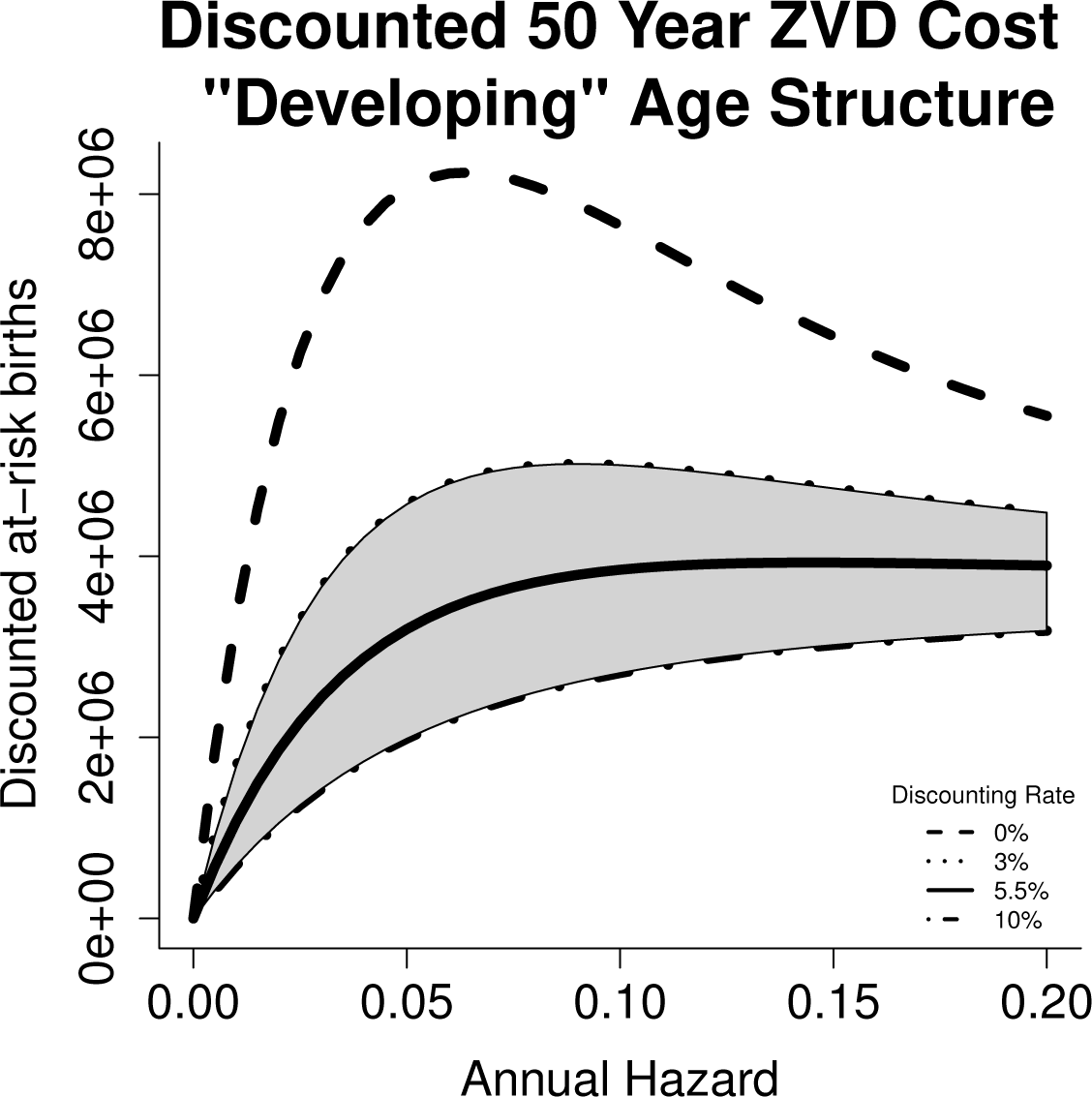
Intermediate levels of ZVD hazard lead to the largest number of at-risk births. When annual discounting of 5.5% or greater is applied to future cases, the total weighted cost of ZVD increases monotonically with annual hazard.

**Fig 8.**
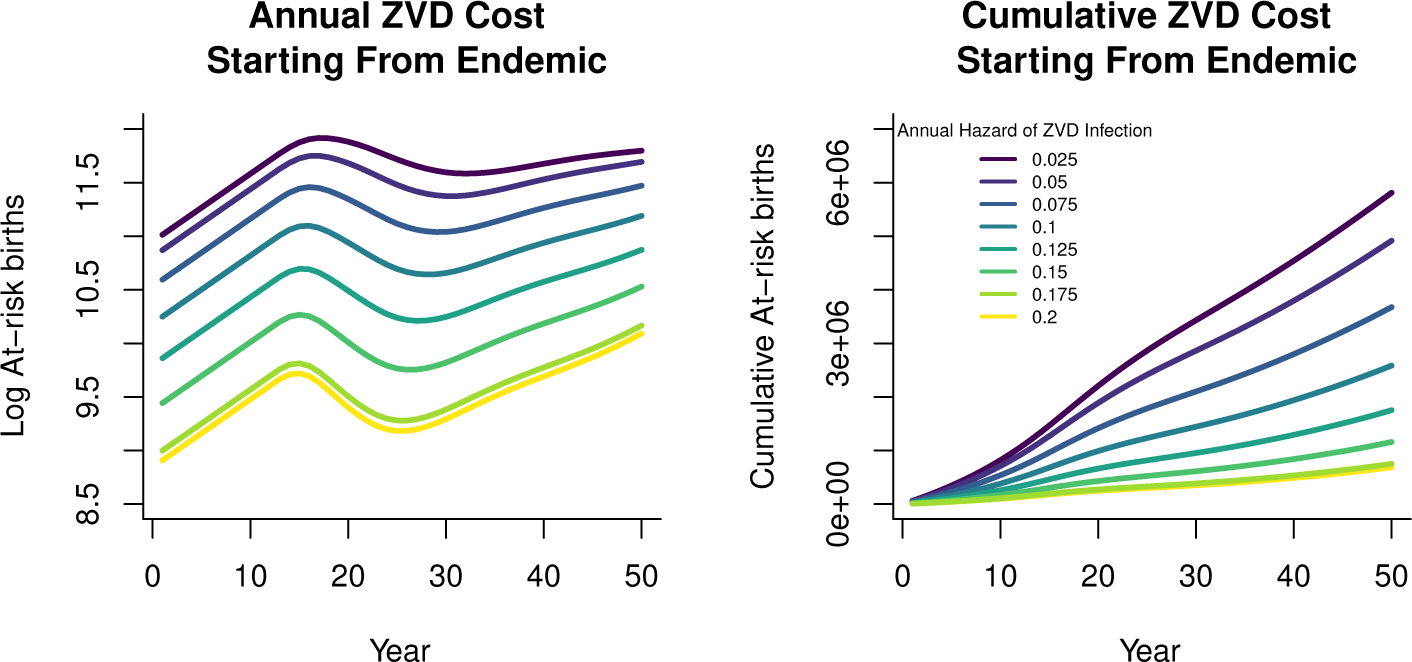
Since the population in the endemic setting begins with partial immunity, annual disease burden is already at equilibrium and fluctuations are primarily due to demographic rates. A clear paradox of control appears in which reductions in attack rate monotonically increase the mean age of infection and therefore relative burden in at-risk age classes.

**Fig 9.**
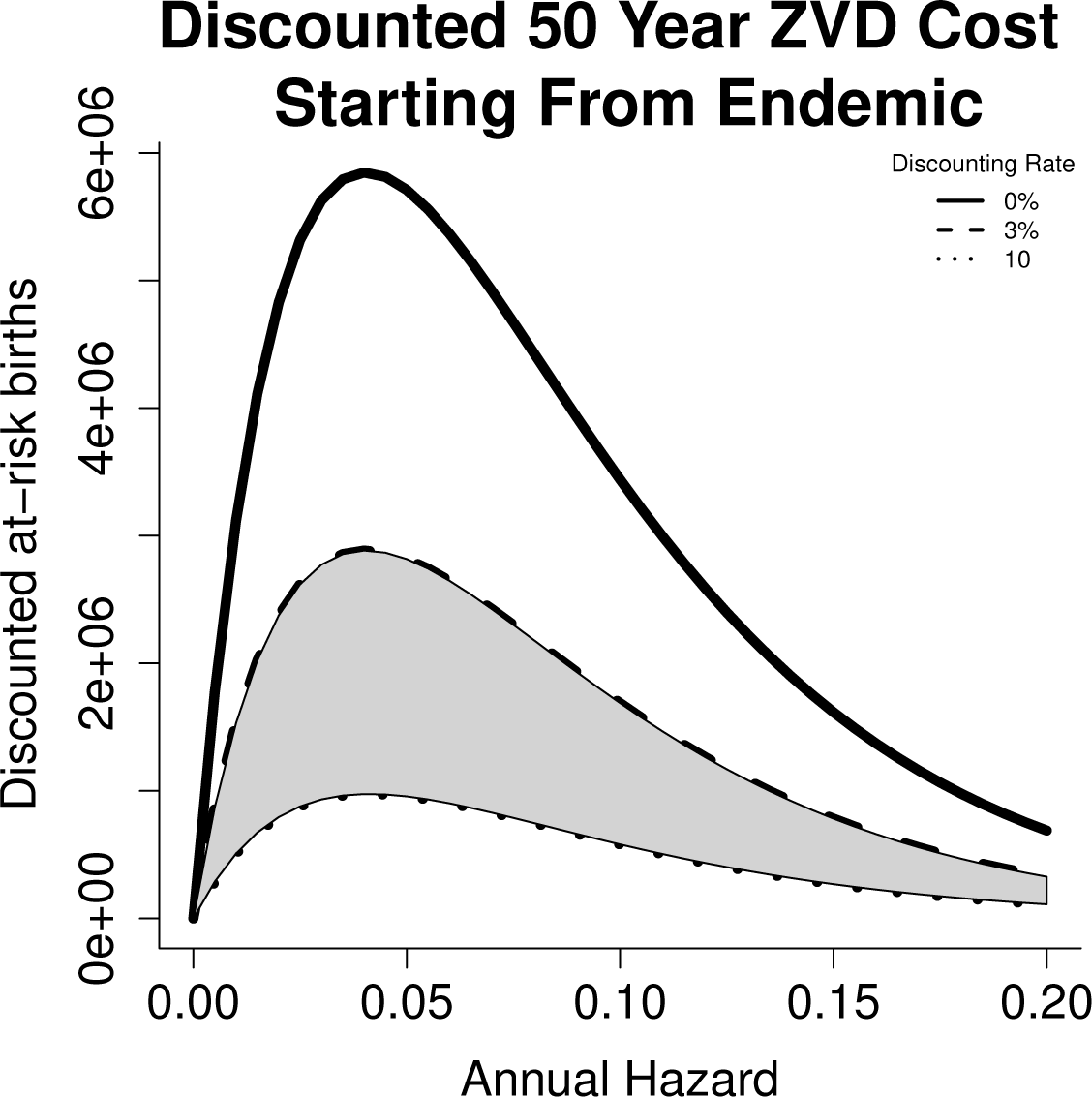
The paradox of control seen in the endemic setting is a function of equilibrium disease burden, and therefore insensitive to discounting. For illustrative purposes we include the range of common discounting rates in the gray shaded region.

**Fig 10.**
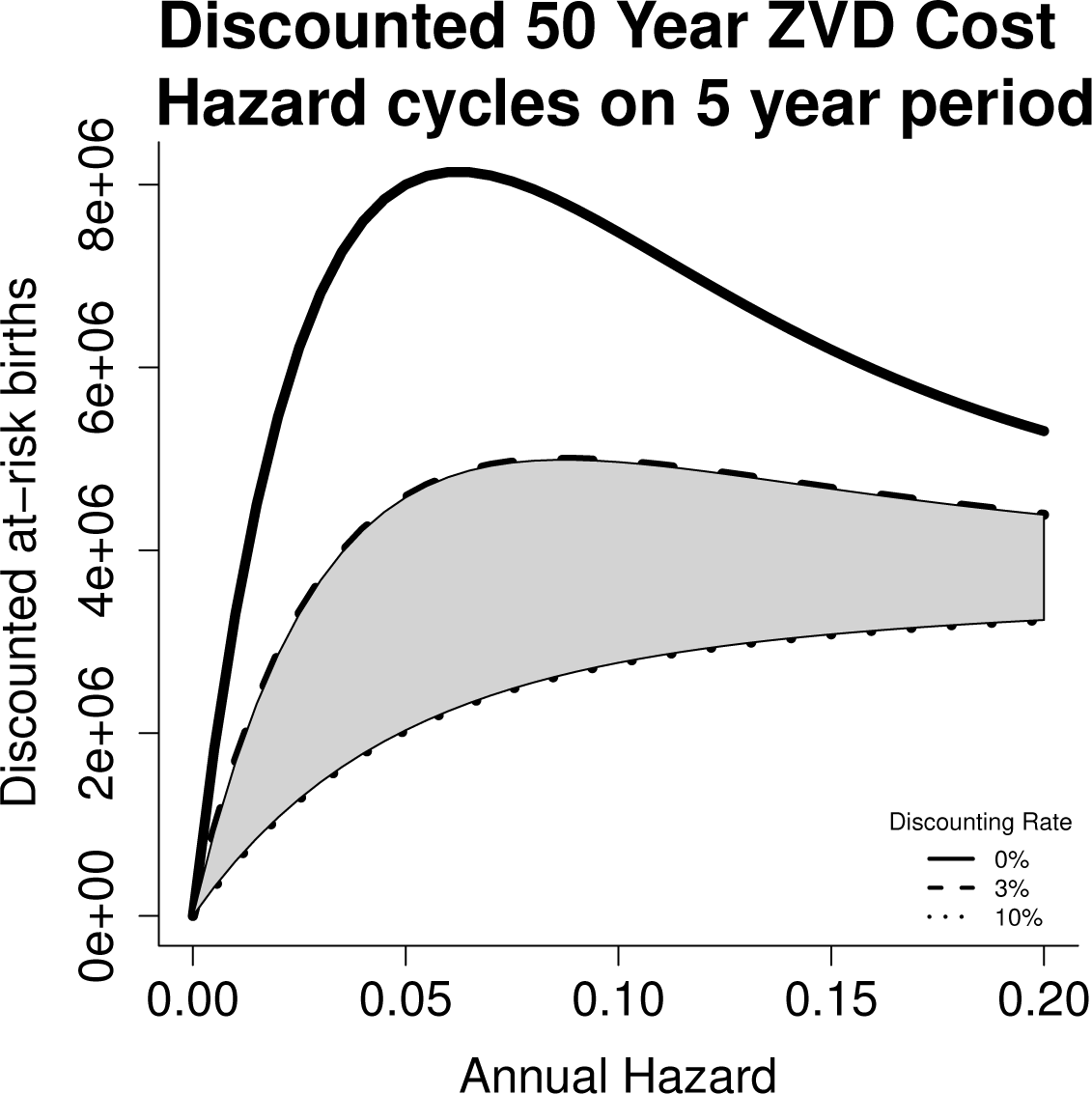
In case of cycles in hazard rates, our qualitative conclusions are broadly unchanged. However, sufficiently long cycles may increase the minimum discounting rate necessary to eliminate the paradox of control by permitting cohorts born during the lower-hazard phase of the cycle to reach reproductive age before encountering ZVD infection. As an illustrative example, we consider below hazard rates that cycle on a five year period.

However, most of the additional cases will occur later in time, by which point there may be medical advances in prenatal care or ZVD control that mitigate the potential for harm. To account for potential discounting of distant future cases relative to near future cases, we weight ZVD cases in our simulation according to the geometric rate 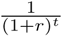 for cases *t* years in the future and a discounting rate *r.* When penalizing current cases more heavily than future cases, we find a reversal of our initial result, returning to the intuitive conclusion that more zika is always worse than less. In the case of our projection based on Brazilian demographics, any discounting rate greater than 1.1% is sufficient to eliminate the paradox of control. When considering potential demographic regimes which differ from those currently prevailing in Brazil (with a disproportionate density of the population in reproductive age classes) we require a higher discounting rate in order to eliminate the possibility of a paradox of control,S1 but the required discount rate is still within typical ranges used in setting social policy.

## Discussion

Higher transmission intensities effectively frontload the total burden of ZVD incidence. Some have argued [32] that the higher long-term incidence of ZVD in low transmission settings should be considered a point against aggressive control efforts to reduce ZIKV transmission. However, policymakers may not be neutral in regards to the timing of potential cases. If two scenarios project similar numbers of cases, it may be preferable to follow the one that delays the burden until later years in light of the expectation that new treatments or preventive measures may be developed in the meantime. Likewise, even a scenario that predicts more total cases over a long time window may be preferred if it involves a lower level of incidence over the time frame of greatest interest to decision makers. While we affirm the central finding of Bewick et al that efforts to limit the size of an initial outbreak of an introduced pathogen such as ZIKV must be traded off against the implications for population-level immunity and long term incidence, we emphasize that possible future cases are less immediately pressing than current cases, and factoring in this prioritization of the present reinforces the importance of limiting disease exposure for at-risk individuals.

We wish to emphasize that our results do not contradict the well-established concept of endemic stability used to justify the avoidance of rubella immunization in some countries. Our conclusions that discounted cumulative future costs from ZIKV are effectively monotonically increasing in infection risk only applies at the start of an epidemic when the population is entirely naive. As the population ages and the infection incidence approaches endemic equilibrium levels, the paradox of control re-emerges (see supplement S2).

Our model does not account for all possible details of long-term ZVD dynamics - the true picture is likely more complicated due to the uncertainty about the extent of sexual transmission [33, 34] and similarity to dengue virus transmission [35]. To the extent that ZVD outcomes depend on the stage of pregnancy and mosquito population density aligns (or not) with human birth seasonality [6, 36], our projections may overstate the total ZVD burden. However, it does so in a uniform way, without biasing comparisons of different transmission intensities. We do not account for costs of ZVD aside from transmission, such as potential strain interactions with dengue fever [37] or link to Guillain-Barre Syndrome [38], both of which would increase the accounting of near-term costs and decrease the future preference for ZVD infection in early childhood.

## Supporting Information

### S1 Under Alternate Demographic Regimes

We consider also a stylized “developing world” age distribution, with a heavily child-biased age distribution. In this case, the relative costs of near-term and future cases are shifted by the smaller proportion of the population currently at risk, and a steeper discounting rate of 5.55% is necessary to eliminate the paradox of control.

### S2 An Endemic Setting

To illustrate the divergence from previous literature’s finding of a paradox of control in rubella-endemic settings, we consider the same projections but with a population whose initial susceptible age structure corresponds to having had a constant infection risk over their lifetimes. This approximates an endemic-disease scenario. In this endemic context, a paradox of control materializes for all discounting rates because there is no initial large outbreak in the higher-hazard scenarios to offset lower long-term caseload in reproductive individuals.

### S3 Time-Varying Hazard Rates

In case of cycles in hazard rates, our qualitative conclusions are broadly unchanged. However, sufficiently long cycles may increase the minimum discounting rate necessary to eliminate the paradox of control by permitting cohorts born during the lower-hazard phase of the cycle to reach reproductive age before encountering ZVD infection. As an illustrative example, we consider below hazard rates that cycle on a five year period.

## References

1. Dick GWA, Kitchen SF, Haddow AJ. Zika Virus (I). Isolations and serological specificity. Trans R Soc Trop Med Hyg. 1952;46(5):509–520. doi:10.1016/0035-9203(52)90042-4.

2. Weaver SC, Costa F, Garcia-Blanco MA, Ko AI, Ribeiro GS, Saade G, et al. Zika virus: History, emergence, biology, and prospects for control. Antiviral Research. 2016;130:69–80. doi:10.1016/j.antiviral.2016.03.010.

3. Costa e Castro M, Álvares da Silva JA, o Nardi ACF, Beltrame A, de Azeredo Costa E, dos Santos L. Protocolo de vigilância e resposta à ocorrência de microcefalia, Versão 1.3. Ministério da Saúde; 2016.

4. Mlakar J, Korva M, Tul N, Popović M, Poljšak-Prijatelj M, Mraz J, et al. Zika Virus Associated with Microcephaly. New England Journal of Medicine. 2016;374(10):951–958. doi:10.1056/NEJMoa1600651.

5. Gao D, Lou Y, He D, Porco TC, Kuang Y, Chowell G, et al. Prevention and control of Zika fever as a mosquito-borne and sexually transmitted disease. arXiv:160404008 [q-bio]. 2016;.

6. Martinez ME. Preventing Zika Virus Infection during Pregnancy Using a Seasonal Window of Opportunity for Conception. PLOS Biology. 2016;14(7):e1002520. doi:10.1371/journal.pbio.1002520.

7. Woog V, Singh S, Browne A, Philbin J. Adolescent Women’s Need for and Use of Sexual and Reproductive Health Services in Developing Countries. Guttmacher Institute; 2015. Available from: https://www.guttmacher.org/sites/default/files/pdfs/pubs/Adolescent-SRHS-Need-Developing-Countries.pdf.

8. Benelli G, Mehlhorn H. Declining malaria, rising of dengue and Zika virus: insights for mosquito vector control. Parasitol Res. 2016; p. 1–8. doi:10.1007/s00436-016-4971-z.

9. Pitzer VE, Lipsitch M. Exploring the relationship between incidence and the average age of infection during seasonal epidemics. J Theor Biol. 2009;260(2):175–185. doi:10.1016/j.jtbi.2009.06.008.

10. Fefferman NH, Naumova EN. Dangers of vaccine refusal near the herd immunity threshold: a modelling study. The Lancet Infectious Diseases. 2015;15(8):922–926. doi:10.1016/S1473-3099(15) 00053-5.

11. Cooper LZ. The History and Medical Consequences of Rubella. Rev Infect Dis. 1985;7(Supplement_1):S2–S10. doi: 10.1093/clinids/7. Supplement 1.S2.

12. Miller E, Gay N. Effect of Age on Outcome and Epidemiology of Infectious Diseases. Biologicals. 1997;25(2):137–142. doi:10.1006/biol.1997.0072.

13. Wesolowski A, Mensah K, Brook CE, Andrianjafimasy M, Winter A, Buckee CO, et al. Introduction of rubella-containing-vaccine to Madagascar: implications for roll-out and local elimination. Journal of The Royal Society Interface. 2016;13(117):20151101. doi:10.1098/rsif.2015.1101.

14. Coleman PG, Perry BD, Woolhouse MEJ. Endemic stability – a veterinary idea applied to human public health. Lancet. 2001;357:1284–1286.

15. Reluga TC, Medlock J, Poolman E, Galvani AP. Optimal Timing of Disease Transmission in an Age-Structured Population. Bull Math Biol. 2007;69(8):2711–2722. doi:10.1007/s11538-007-9238-5.

16. Reluga TC, Li J. Games of age-dependent prevention of chronic infections by social distancing. Journal of Mathematical Biology. 2012;66(7):1527–1553. doi:10.1007/s00285-012-0543-8.

17. Panagiotopoulos T, Berger A, Antoniadou L, Valassi-Adam E. Increase in congenital rubella occurrence after immunisation in Greece: retrospective survey and systematic review: How does herd immunity work? BMJ. 1999;319(7223):1462–1467. doi:10.1136/bmj.319.7223.1462.

18. Vynnycky E, Gay NJ, Cutts FT. The predicted impact of private sector MMR vaccination on the burden of Congenital Rubella Syndrome. Vaccine. 2003;21(21–22):2708–2719. doi:10.1016/S0264-410X(03)00229-9.

19. Yakob L, Walker T. Zika virus outbreak in the Americas: the need for novel mosquito control methods. The Lancet Global Health. 2016;4(3):e148–e149. doi:10.1016/S2214-109X(16) 00048-6.

20. Cohen J. The race for a Zika vaccine is on. Science. 2016;351(6273):543–544. doi:10.1126/science.351.6273.543.

21. Tripp RA, Ross TM. Development of a Zika vaccine. Expert Review of Vaccines. 2016;15(9):1083–1085. doi:10.1080/14760584.2016.1192474.

22. Marston HD, Lurie N, Borio LL, Fauci AS. Considerations for Developing a Zika Virus Vaccine. New England Journal of Medicine. 2016;375(13):1209–1212. doi:10.1056/NEJMp1607762.

23. Harrison M. Valuing the Future: the social discount rate in cost-benefit analysis. Australian Government Productivity Commission; 2010. Available from: http://www.pc.gov.au/research/supporting/cost-benefit-discount.

24. United Nations DoE, Social Affairs PD. World Fertility Data 2012. United Nations; 2013. Available from: http://data.un.org/DocumentData.aspx?id=319.

25. Honein MA, Dawson AL, Petersen EE, Jones AM, Lee EH, Yazdy MM, et al. Birth Defects Among Fetuses and Infants of US Women With Evidence of Possible Zika Virus Infection During Pregnancy. JAMA. 2016; doi:10.1001/jama.2016.19006.

26. Brasil P, Pereira JPJ, Moreira ME, Ribeiro Nogueira RM, Damasceno L, Wakimoto M, et al. Zika Virus Infection in Pregnant Women in Rio de Janeiro. New England Journal of Medicine. 2016;375(24):2321–2334. doi:10.1056/NEJMoa1602412.

27. Cauchemez S, Besnard M, Bompard P, Dub T, Guillemette-Artur P, Eyrolle-Guignot D, et al. Association between Zika virus and microcephaly in French Polynesia, 2013–15: a retrospective study. The Lancet. 2016;387(10033):2125–2132. doi:10.1016/S0140-6736(16)00651-6.

28. Kermack WO, McKendrick AG. A contribution to the mathematical theory of epidemics. Proceedings of the Royal Society A. 1927;115:700–721. doi:10.1098/rspa.1927.0118.

29. Grenfell BT, Anderson RM. Pertussis in England and Wales: An Investigation of Transmission Dynamics and Control by Mass Vaccination. Proceedings of the Royal Society B: Biological Sciences. 1989;236(1284):213–252. doi:10.1098/rspb.1989.0022.

30. McKendrick AG. Applications of mathematics to medical problems. Proceedings of the Edinburgh Mathematical Society. 1926;44:98–130. doi:10.1017/s0013091500034428.

31. Lessler J, Metcalf CJE. Balancing Evidence and Uncertainty when Considering Rubella Vaccine Introduction. PLOS ONE. 2013;8(7):e67639. doi:10.1371/journal.pone.0067639.

32. Bewick S, Fagan WF, Calabrese JM, Agusto F. Zika Virus: Endemic Versus Epidemic Dynamics and Implications for Disease Spread in the Americas. bioRxiv. 2016; p. 041897. doi:10.1101/041897.

33. Musso D, Roche C, Robin E, Nhan T, Teissier A, Cao-Lormeau VM. Potential Sexual Transmission of Zika Virus. Emerging Infectious Diseases. 2015;21(2):359–361. doi:10.3201/eid2102.141363.

34. Harrower J, Kiedrzynski T, Baker S, Upton A, Rahnama F, Sherwood J, et al. Sexual Transmission of Zika Virus and Persistence in Semen, New Zealand, 2016. Emerging Infectious Diseases. 2016;22(10):1855–1857. doi:10.3201/eid2210.160951.

35. Carlson C, Dougherty E, Getz W. An ecological assessment of the pandemic threat of Zika virus. bioRxiv. 2016; p. 040386. doi:10.1101/040386.

36. Martinez-Bakker M, Bakker KM, King AA, Rohani P. Human birth seasonality: latitudinal gradient and interplay with childhood disease dynamics. Proc R Soc B. 2014;281(1783):20132438. doi:10.1098/rspb.2013.2438.

37. Dejnirattisai W, Supasa P, Wongwiwat W, Rouvinski A, Barba-Spaeth G, Duangchinda T, et al. Dengue virus sero-cross-reactivity drives antibody-dependent enhancement of infection with zika virus. Nat Immunol. 2016;17(9):1102–1108. doi:10.1038/ni.3515.

38. Parra B, Lizarazo J, Jiménez-Arango JA, Zea-Vera AF, González-Manrique G, Vargas J, et al. Guillain–Barré Syndrome Associated with Zika Virus Infection in Colombia. New England Journal of Medicine. 2016;375(16):1513–1523. doi:10.1056/NEJMoa1605564.

